# 100 Million Years of Solitude: Tracing the Origins and Diversification of AZG Cytokinin Transporters

**DOI:** 10.64898/2026.07.17.739114

**Authors:** Maximilian Klamke, Veronica G. Maurino, Christopher Grefen, Marcelo Desimone, Tomas M. Tessi

## Abstract

**Dear editor:** The evolution of complex plant body architectures required the refinement of hormone transport networks, yet the evolutionary origins and functional diversification of cytokinin transporters remain unclear. Our understanding of cytokinin (CK) transport across cellular membranes has advanced substantially (Tessi and Maurino, 2024). Among CK transport systems, the AZA-GUANINE RESISTANT (AZG) transporter family has attracted particular interest (Tessi et al., 2020; Tessi et al., 2023). The determination of AZG1 and AZG2 structures has revealed their transport mechanisms, making AZGs among the best-characterized phytohormone transporters (Xu et al., 2024; Wei et al., 2025). Here, we reconstruct the molecular evolution of the AZG family and show that AZG1 represents the ancestral land plant orthologue, preserving conserved proton-coupling residues found in streptophyte algae. Conversely, AZG2 likely emerged during vascular plant diversification and displays relaxed evolutionary constraints within its ligand-binding pocket, providing a molecular basis for a shift toward proton independence. Structural modeling and split-ubiquitin assays further reveal that despite this divergence, AZG1 and AZG2 retain the capacity to heterodimerize. Together, our findings uncover the stepwise innovation of the AZG family and link structural diversification to the increasing complexity of plant hormone transport networks.

---

The AZG protein family phylogeny resolves a monophyletic bacterial outgroup that is included within the AZG-like group (Fig. 1A). Algal AZG-like sequences branch basal to fungal and streptophyte clades, indicating retention of an ancestral AZG-like orthologue across green algae. The fungal-specific AZGA clade is sister to Streptophyta AZGs, consistent with an early independent divergence of the fungal lineage. Notably, some Charophyta taxa contain a second AZG sequence. This variant occupies the earliest-diverging position within the streptophyte AZG clade, suggesting that the sequence divergence gives rise to embryophyte AZGs predating land plant terrestrialization. This is consistent with the hypothesis that rudimentary cytokinin signaling networks were already present in charophyte algae (Powell and Heyl, 2023).

**Figure 1.**
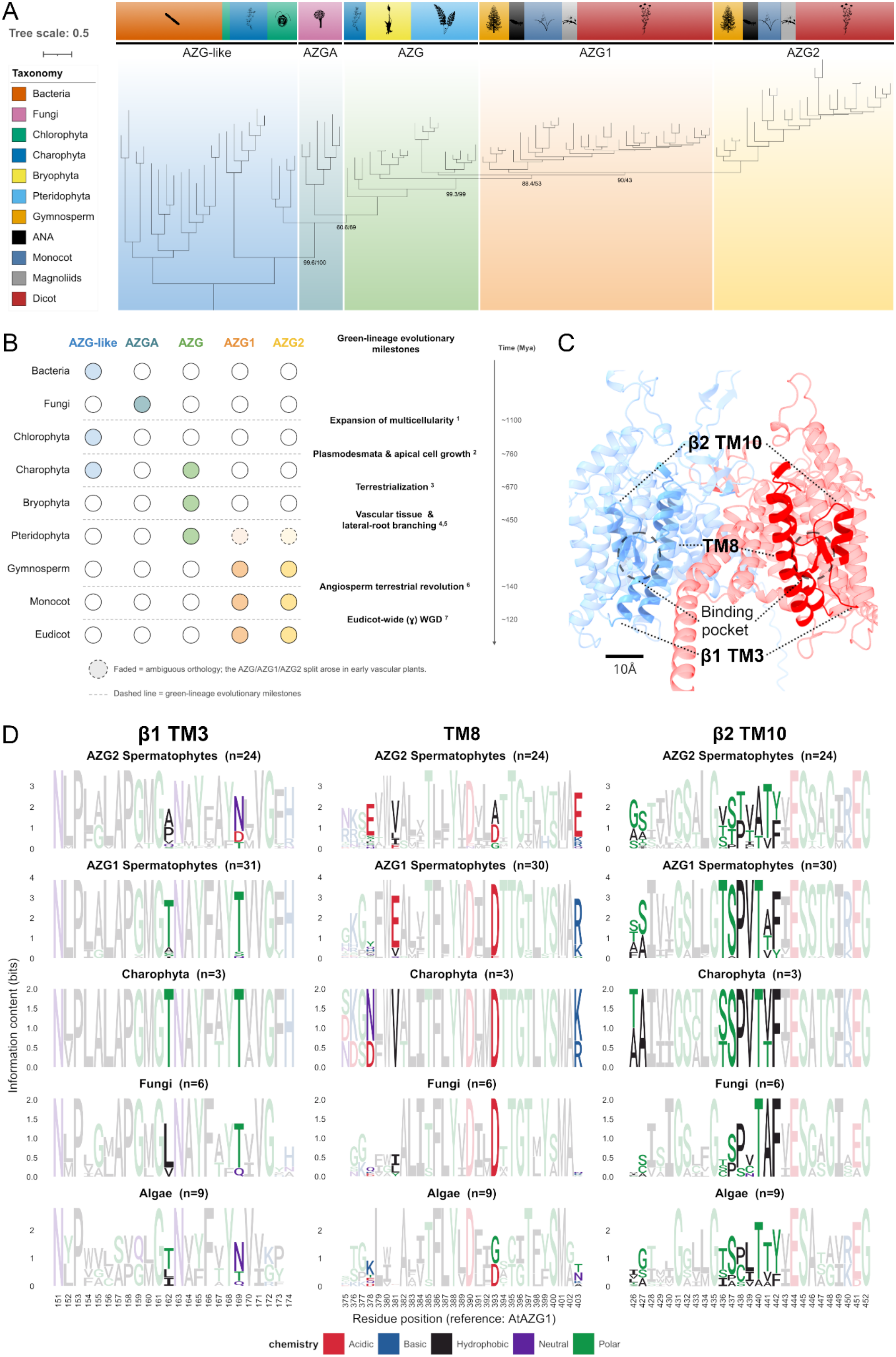
Phylogenetic tree and molecular evolution of the AZG transporter family. **(A)** AZG-like sequences of representative species from a selective proteome data set were identified via BLASTp (Pucker and Iorizzo, 2023) against AtAZG1 (AT3G10960), AtAZG2 (AT5G50300), and AZGA of *Aspergillus nidulans*. Sequences were aligned using MAFFT L-INS-i (Katoh and Standley, 2013) and poorly aligned regions were trimmed using the kpic-smart-gap algorithm of ClipKIT (Steenwyk et al., 2020). Scaffold-derived or long-branched sequences, and species-specific duplicates were excluded to improve tree resolution and computational efficiency. Lycophyte sequences (*Selaginella moellendorffii* and *Diphasiastrum complanatum*) were additionally excluded, as they failed to form a monophyletic group and instead resolved with unusually long branches that destabilized overall tree topology. The phylogenetic tree was inferred with IQ-TREE (Nguyen et al., 2015) using the Q.PFAM+I+R7 model, 1000 ultrafast bootstrap (UFBoot) replicates and 1000 SH-aLRT replicates. Displayed node labels indicate SH-aLRT/UFBoot support values at key splits; basal nodes are unlabeled due to midpoint rooting. Tree visualization and midpoint rooting were performed in iTOL v6 (Letunic and Bork, 2024). For clarity, node support values and sequence headers are not shown. The original tree file, species dataset, and all excluded sequences are available in Supplementary Data S1. Lineage silhouettes were obtained from https://www.phylopic.org/. **(B)** Simplified schematic representation of the evolutionary distribution and expansion of AZG-like transporters across major lineages from bacteria and fungi to green plants. Colored circles indicate the presence of the corresponding AZG ortholog; uncolored circles indicate its absence. Dashed lines denote key evolutionary milestones within Archaeplastida; numbered footnotes (1–7) indicate the corresponding sources: ^1^Bowles et al. (2023), ^2^Nishiyama et al. (2018), ^3^Harris et al. (2022), ^4^Woudenberg et al. (2022), ^5^Hetherington et al. (2020), ^6^Benton et al. (2022), and ^7^Aköz and Nordborg (2019). A vertical time scale provides approximate evolutionary context across these milestones. Faded elements indicate uncertain orthology relationships associated with the AZG family split within pteridophytes. **(C)** Sequence logos for the β-sheet 1 and transmembrane α-helix 3 (β1–TM3), Transmembrane α-helix 8 (TM8) and β-sheet 2 and Transmembrane α-helix 10 (β2-TM10). The sequence clusters have been determined based on the tree in (A). Regions and sequence positions are in reference to AtAZG1. **(D)** Overview of the AZG1 (blue) AZG2 (red) heterodimer highlighting the regions with high conservational constraints analysed in the sequence logos. The scale represents 10 Å. **Figure 1 Alt-Text**. Phylogenetic and sequence conservation analysis of the AZG transporter family. **A:** Phylogenetic tree and taxonomy distribution chart mapping AZG subfamilies (AZG-like, AZGA, AZG, AZG1, AZG2), from bacteria and fungi to gymnosperms and angiosperms. **B:** Diagram illustrating the evolutionary lineage expansion and duplication history of AZG transporters in land plants. **C**: scheme of AZG1 and AZG2 heterodimer depicting highly conserved domains participating in the transporter binding pocket (β1-TM3, TM8, β2-TM10). **D:** Sequence alignment logos depicting bits of information content and amino acid conservation across key binding pocket regions (β1-TM3, TM8, β2-TM10) among Spermatophytes, Charophyta, Fungi, and Algae.

Within bryophytes, AZG homologues are generally single-copy, whereas pteridophytes demonstrate the first plant-type duplication event. Gymnosperms represent the earliest lineage in which AZGs cleanly resolve into two clades. Across angiosperms, AZGs are retained in all major lineages. Together, these data indicate a stepwise expansion of the family. Plant-type AZGs likely arose via duplication of an ancestral AZG-like gene in charophytes. The retention of a single isoform in bryophytes suggests the loss of additional AZG-like genes at or prior to land plant emergence. The plant-type AZG split arose later within tracheophytes, with duplications in pteridophytes and a clean AZG1/AZG2 separation in seed plants (Fig. 1B). This diversification may have contributed to the emergence of novel hormone transport strategies associated with land plant complexity, including the regulation of lateral root branching.

Based on structural and mechanistic analyses of *A. thaliana* AZG1 and AZG2 (AtAZG) (Xu et al., 2024; Wei et al., 2025), we focused on three regions involved in substrate binding and stabilization within the transporter domain (Fig. 1C-D; Fig. S1). These regions are highly conserved, although variability occurs across AZG orthologues. Within the β1-TM3 region, T162 is conserved in most AZG1 sequences, whereas AZG2 homologues frequently contain non-polar residues, mainly alanine or proline, at this position (Fig. 1D). In AZG1, T162 participates in a proposed hydrogen-bonding network with D390 in TM8 and T440 in β2-TM10 (Xu et al., 2024). In contrast to AZG1, residues T162 and T440 are poorly conserved in AZG2, while D390 (D345 in AZG2) stabilizes the side chain of tZ instead of participating in the H-bond network. The presence of such a conserved network may support proton-coupled transport, and its relaxation in AZG2 may explain differences in proton dependence between both transporters (Tessi et al., 2020; Wei et al., 2025). Charophyte and bryophyte sequences retain the AZG1-type configuration, supporting AZG1 as the ancestral plant AZG from which AZG2 later evolved.

Several residues within the substrate binding pocket, including Y62, M160, D390, and E444, are conserved across AZGs (Fig. 1D). However, structural studies revealed distinct substrate orientations in AtAZG1 and AtAZG2 (Xu et al., 2024; Wei et al., 2025), reflected by divergence at positions 381, 393, 403, 427, 440, 441, and 442. Whereas charophyte, bryophyte, and fern sequences retain an AZG1-like configuration, AZG2 homologues display a distinct pattern consistent with a derived transporter lineage. Several lineage-specific residues surrounding the substrate-binding pocket show substantial physicochemical changes and may contribute to protein interactions or substrate recruitment/release, including AZG1-to-AZG2 substitutions S426→G380, R403→E358, and H378→E333 (Fig. 1D and S1).

Beyond single amino acid substitutions, the most conspicuous structural difference between AtAZG1 and AtAZG2 is the absence of β-sheets 6-9 in AtAZG2 (Wei et al., 2025; Fig. 2A-B). This AZG2-specific feature appears in the ANA-grade angiosperm *Amborella trichopoda* and is exclusively retained across angiosperms (Fig. S2). Although this region contributes to the scaffold domain and dimer formation, its absence does not prevent AZG2 assembly, as both AtAZG1 and AtAZG2 form dimers in cryo-EM structures (Xu et al., 2024; Wei et al., 2025).

**Figure 2.**
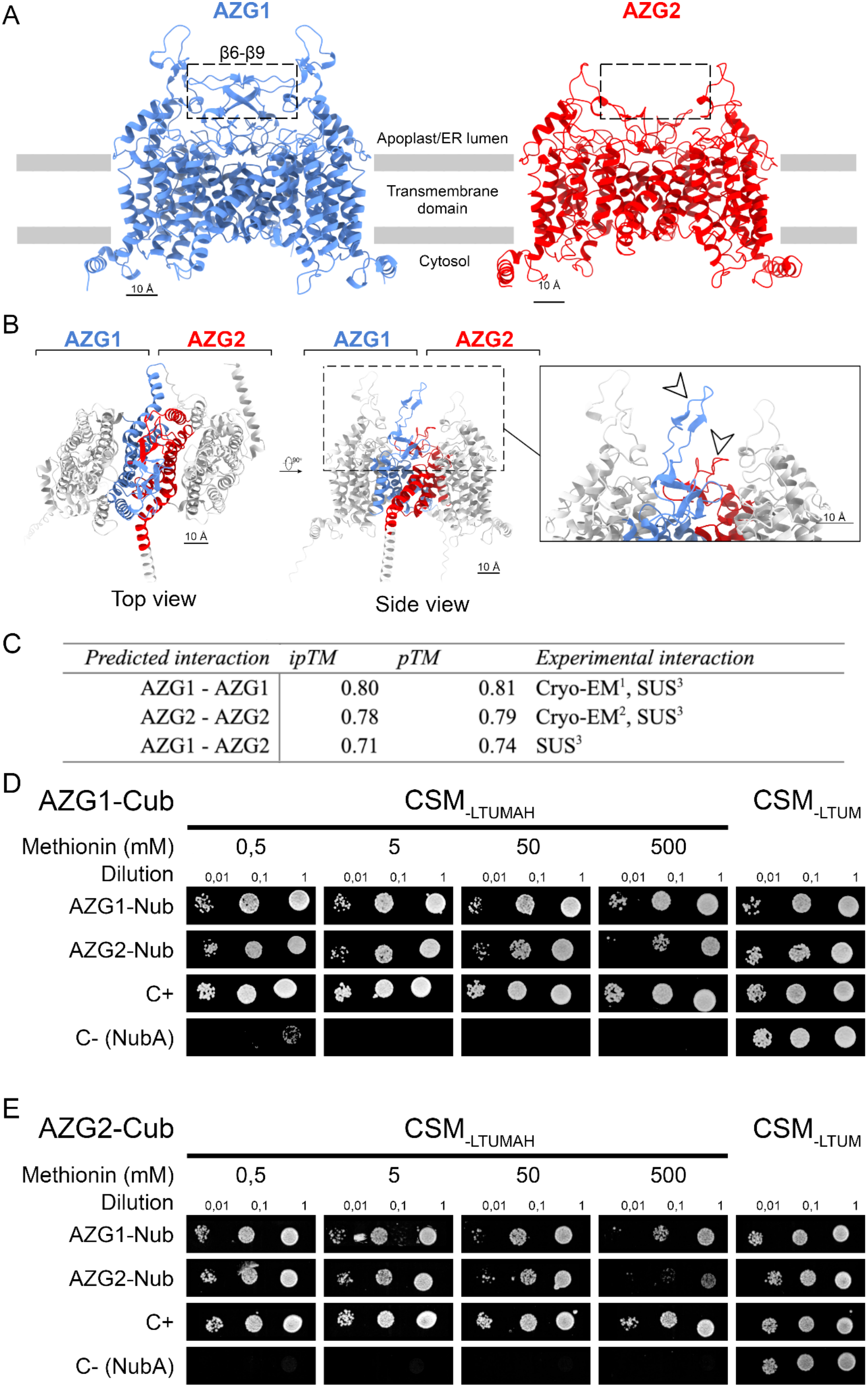
Protein-protein interaction between AZG1-AZG2. **(A)** Comparison of AtAZG1 and AtAZG2 homodimers highlighting the lack of the β6–β9 sheets in AtAZG2. The structure files used for the comparison are publicly available with the identifiers 8WMQ (AZG1) and 9j16 (AZG2) (Xu et al., 2024; Wei et al., 2025). **(B)** AZG1-AZG2 heterodimer structure prediction by AlphaFold 3. Interacting domains are shown in blue for AZG1 and red for AZG2. Arrows point to the extracellular loop involved in the dimer formation showing their different sizes between AZG1 and AZG2. **(C)** AlphaFold 3 folding (pTM) and interaction (ipTM) predictions for *Arabidopsis thaliana* AZGs homo- and heterodimers. Experimental interactions have been reported using the indicated technique in the following works: ^1^ Xu et al., 2024; ^2^ Wei et al., 2025; ^3^ This work; ^4^ Tessi et al., 2023. **(D-E)** Split Ubiquitin System (SUS) interaction test between AZG1 and AZG2. SUS using AZG1 **(D)** or AZG2 **(E)** as Cub versus AZG1 or AZG2 as Nub fusions. Wild-type Ubiquitin was used as a positive control while Nub-A alone was used as a negative control. Selective media (CSM_-LTUMAH_) with increasing concentrations of Methionine were tested to repress the expression of the constructs via the Met25 promoter and regulate the amount of interacting partners. Non-selective media was used to test the viability of the cells (CSM_-LTUM_). **Figure 2 Alt-Text**. Schematic and experimental analysis of AZG1 and AZG2 dimerization. **A–B:** Structural diagrams showing membrane orientations (apoplast/ER lumen, transmembrane domain, cytosol) and interaction interface for AZG1 and AZG2. **C:** Table comparing AlphaFold3 predicted interaction scores (ipTM and PTM) with experimental validation methods (Cryo-EM and split-ubiquitin system) for homodimeric (AZG1-AZG1, AZG2-AZG2) and heterodimeric (AZG1-AZG2) complexes. **D–E:** Split-ubiquitin yeast growth assays showing interaction strength of AZG1-Cub and AZG2-Cub constructs against Nub fusion partners across multiple cell dilutions and methionine concentrations.

AlphaFold 3 predicted AtAZG1 and AtAZG2 homodimers as well as AZG1-AZG2 heterodimers with high confidence (Fig. 2C). Split-Ubiquitin System (SUS) interaction assays confirmed AtAZG1 and AtAZG2 homodimerization and demonstrated *in vivo* reciprocal heterodimerization (Fig. 2D-E; Fig. S3). Under stringent conditions (500 mM methionine), AZG1 homodimers exhibited the strongest interaction, followed by AZG1-AZG2 heterodimers and weaker AZG2 homodimers. This interaction hierarchy aligns with AtAZG1’s larger dimerization interface, indicating greater homodimeric stability. The structural differences between AZG1 and AZG2, particularly at the dimerization interface and transporter vestibule, further suggest that homo- and heteromeric complexes may not be functionally equivalent. Heterodimerization may therefore modulate transport kinetics, substrate preference, proton dependence, and protein turnover, adding a layer of regulatory complexity to coordinated cytokinin transport in the cells where both transporters are co-expressed such as the lateral root overlying tissues.

Together, our evolutionary, structural, and interaction analyses show that AZG diversification involved shifts in gene number, sequence constraints, and oligomeric properties. This oligomeric flexibility may fine-tune cytokinin distribution and coordinate it with auxin transport across tissues, developmental stages, and environments. This work highlights key milestones in AZG evolution and provides a foundation for future experimental studies aimed at elucidating clade-specific neofunctionalization and hormone transport mechanisms within the protein family.

## Acknowledgments

We thank Prof. Dr. Klaus Harter for facilitating the research stay of TT at the ZMBP, Tübingen.

## Author contributions

TT and MD conceived the project. MK, VM, and TT worked on the phylogenetic analysis. TT, MD, and CG designed and carried out the split-ubiquitin experiments. TT performed the structural and protein sequence analysis. All authors contributed to writing the manuscript and approved the submitted version.

## Supplementary data

Supplementary material is available at Plant Physiology online.

## Data availability

The data and methods that support the findings of this study are available in the supplementary data of this article.

## Supplemental Figures

**Suppl. Figure S1.**
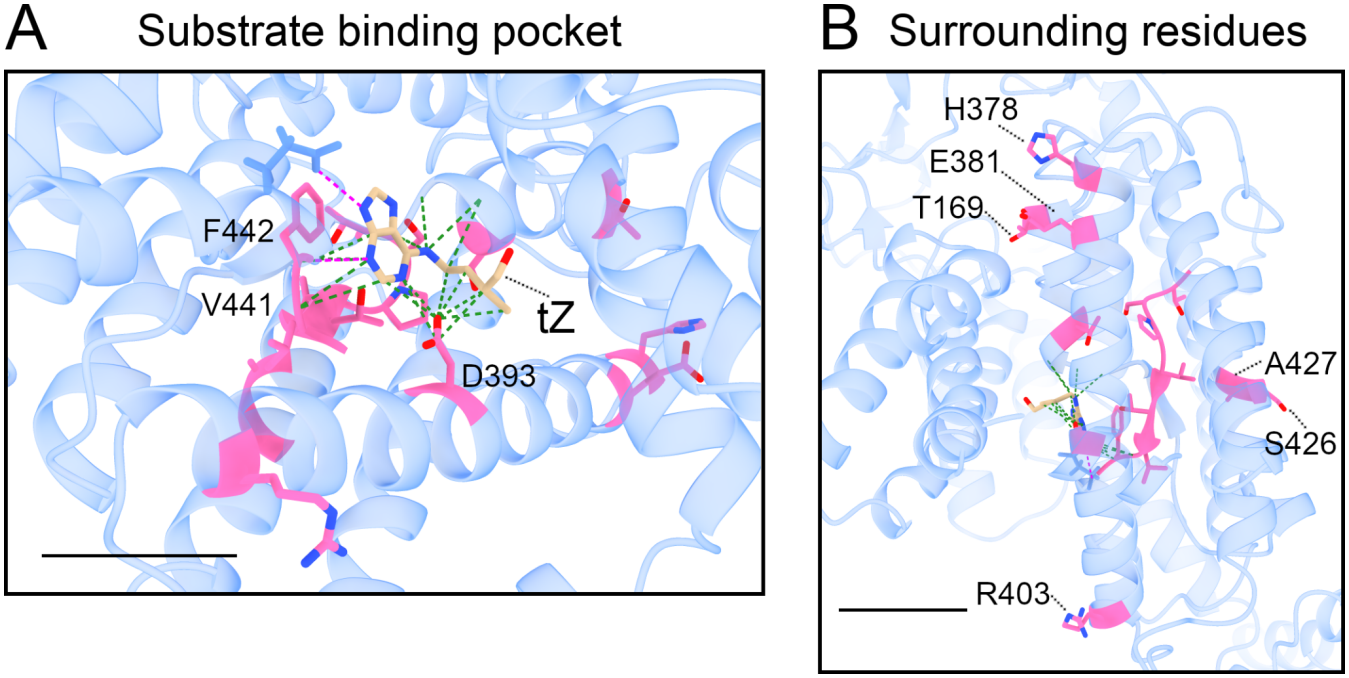
Key residues participating in the stabilization of the transporter substrate. Conserved positions within sequence clusters that are not conserved across different AZG family members are indicated in magenta. In **(A)** are shown the contact points of such residues located in the AZG1 trans-Zeatin (tZ) binding pocket while **(B)** shows the residues that are outside of the pocket and not directly interacting with the substrate. The scale represents 10 Å.

**Suppl. Figure S2.**
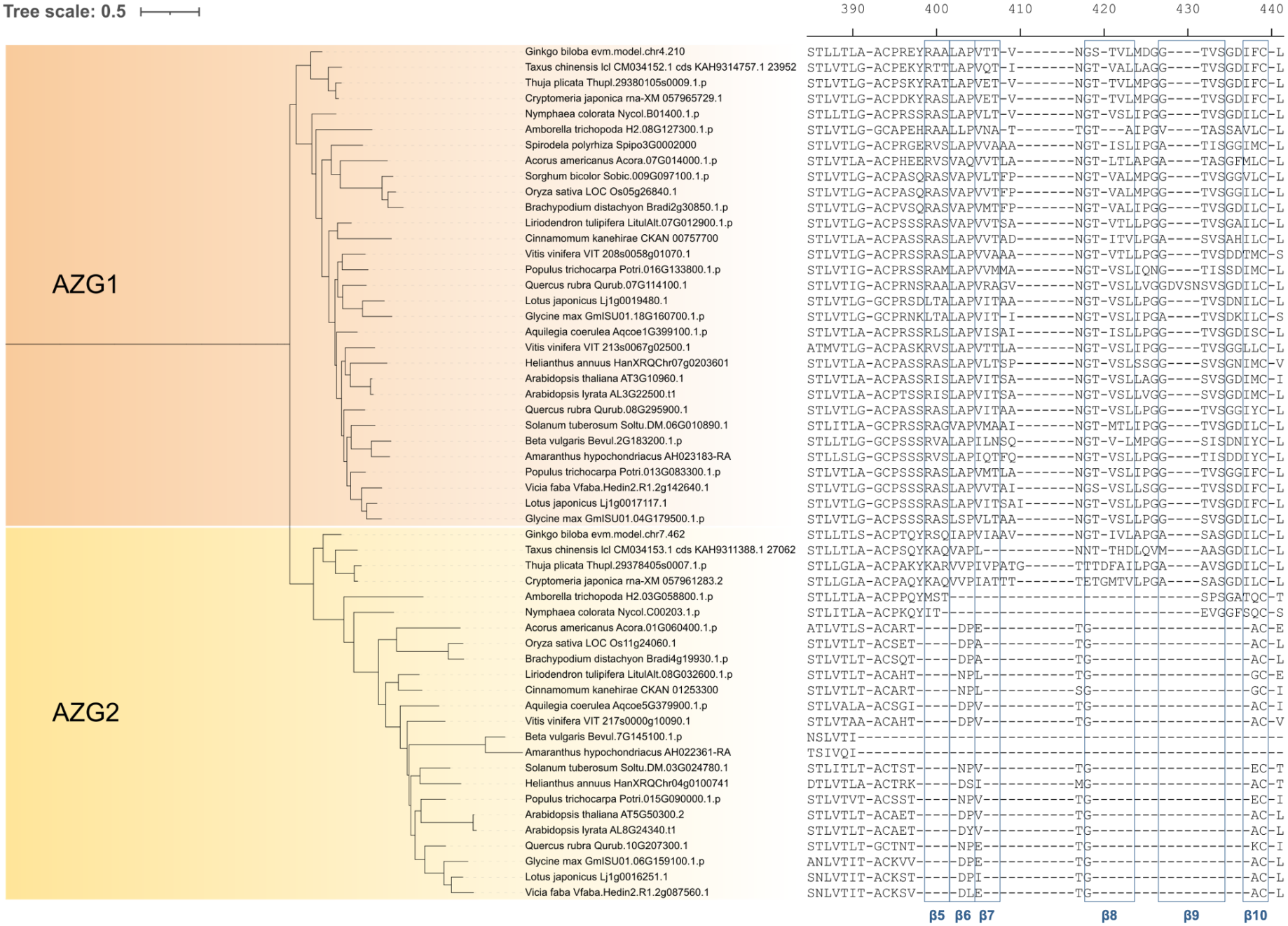
AZG1/AZG2 phylogeny with mapped β-sheets. Pruned phylogenetic tree of AZG-family transporters, showing AZG1 and AZG2 across spermatophytes, displayed alongside the corresponding multiple sequence alignment (MSA). Blue boxes denote β-strands β5–β10, assigned from the MSA based on AtAZG1 following Xu et al. (2024). Within the AZG2 clade, β5 appears shifted to an intermediate position between β6 and β7, and the conserved glycine of β10 is likewise shifted to the first position of β8. As the underlying strand sequences are identical, these apparent positional shifts most likely reflect an artifact caused by the larger sequence number in this MSA relative to the Xu et al. reference alignment.

**Suppl. Figure S3.**
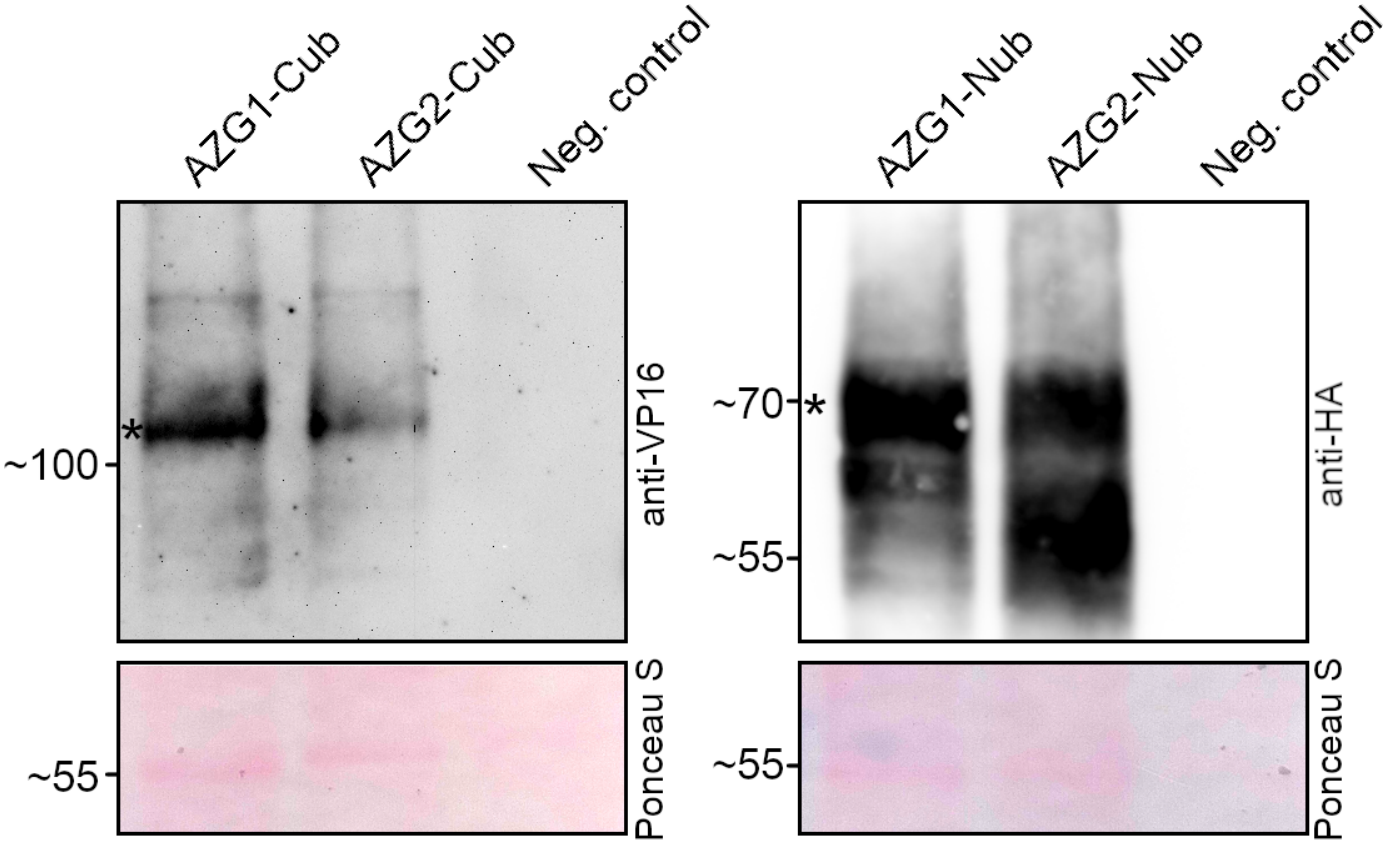
Western blot confirmation of Cub and Nub expression in yeast cultures used for SUS. anti-VP16 was used to detect Cub fusions and anti-HA to detect Nub fusions. Asterisks showed the band corresponding to the Cub- and Nub-fusions. A non-transformed yeast total protein extract was used as a blotting negative control. Total protein loading control was stained using Punceau S.

