## Supplementary figures and images for "100 Million Years of Solitude: Tracing the Origins and Diversification of AZG Cytokinin Transporters"

### AZG_ufbs_basic_inverted.png

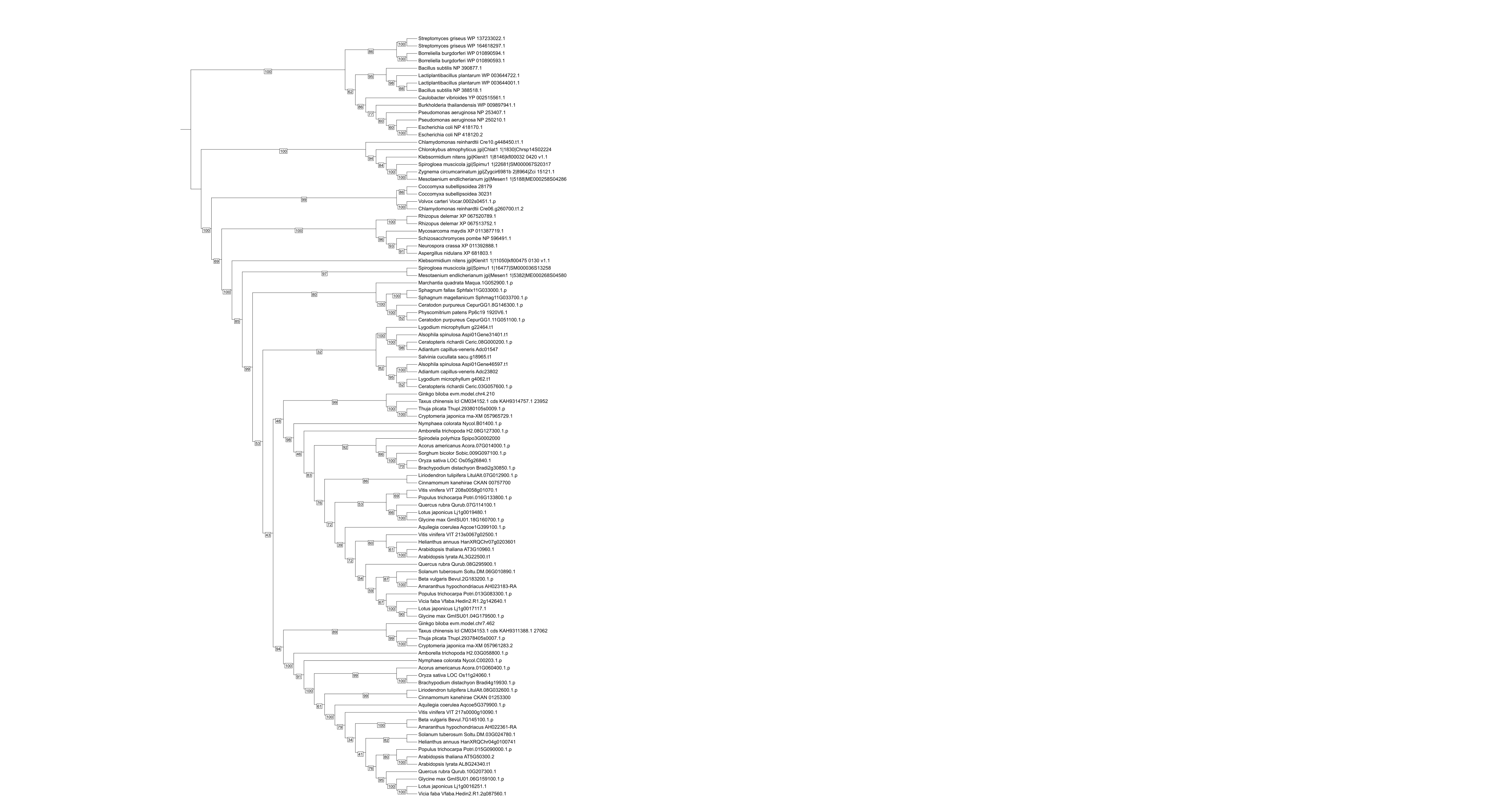

### Fig_S1.png

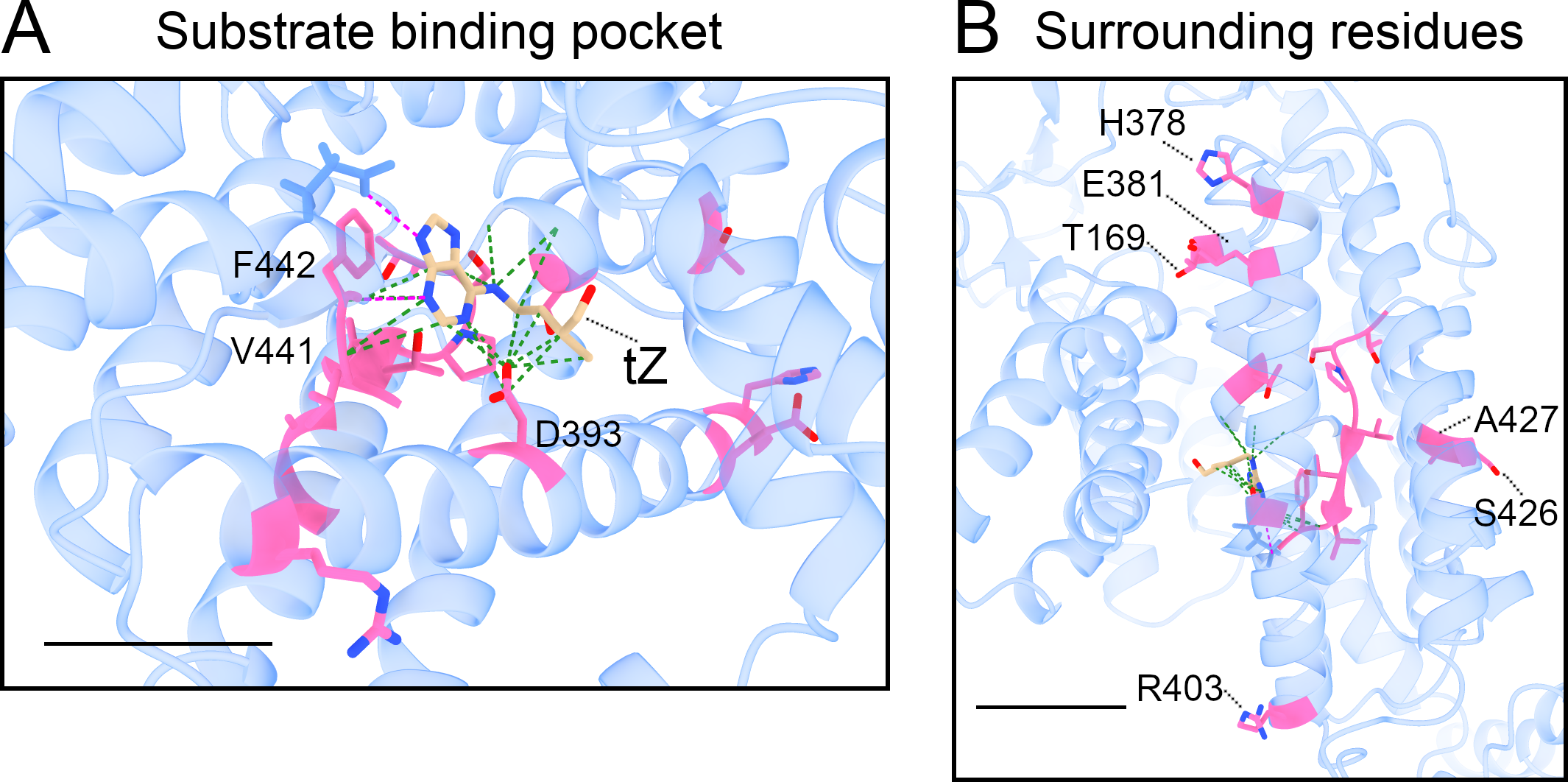

### Fig_S2.png

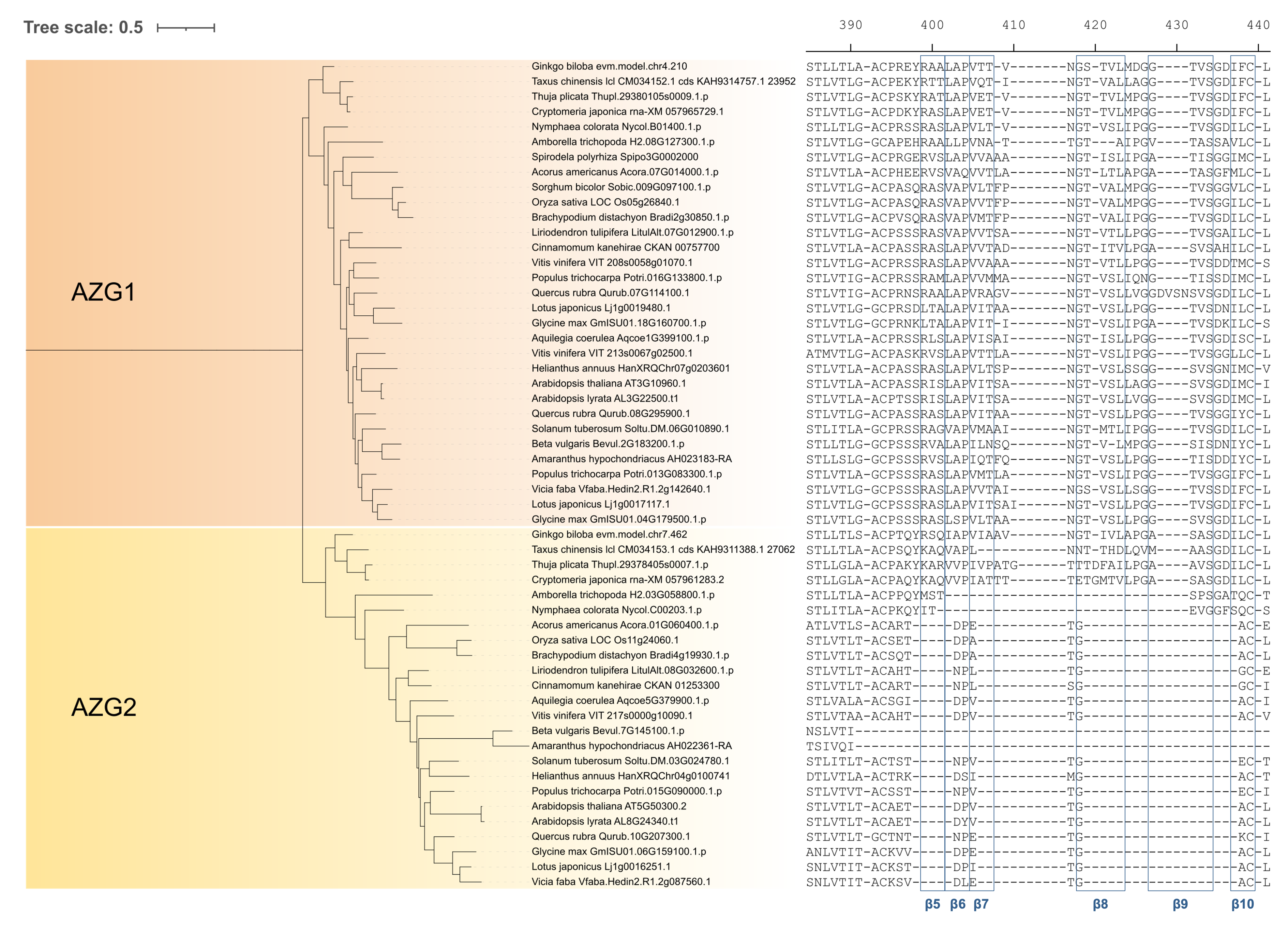

### Fig_S3.png

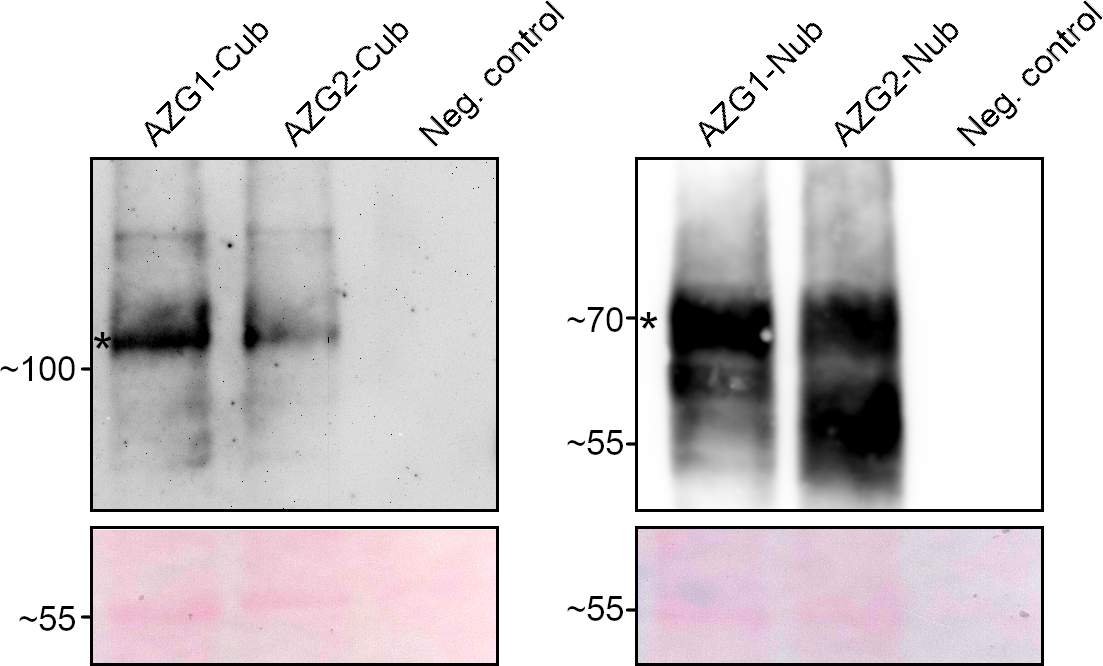
